# Reconstruction of functional olfactory sensory tissue from embryonic nasal stem cells

**DOI:** 10.1101/2024.09.13.612986

**Authors:** Kazuya Suzuki, Fumi Wagai, Mototsugu Eiraku

**Author notes:** To whom correspondence should be addressed: Mototsugu Eiraku, Ph.D., Laboratory of Developmental Systems Institute for Life and Medical Sciences, Kyoto University, Shogoin-Kawahara-cho, Kyoto, 606-8507, Japan.

## Abstract

During the development of the olfactory epithelium (OE), olfactory sensory neurons (OSNs) express only one member of the odorant receptor (OR) gene family, and OSNs expressing the same OR converge their axons to the same set of glomeruli on the olfactory bulb (OB). The resulting odor maps allow mice to discriminate more than 100,000 different odorants using about 1,000 ORs. It remains elusive how odor maps are formed. Here, we show a means of forming OE organoids with pseudostratified structure from mouse embryonic OE stem cells. Single-cell RNA sequencing revealed that the OE organoids give rise to all the OE cellular lineages and undergo active neurogenesis. We also found that most OSNs in OE organoids exclusively express only one type of ORs and exhibit a unique molecular code of axon guidance-related genes that can discriminate between OR classes. Thus, OE organoids could be a useful model for studying olfactory nervous system development.

## Introduction

The olfactory epithelium (OE) is a pseudostratified epithelium derived from the olfactory placode and is composed of cells with different functions. (Cuschieri et al., 1975; Sokpor et al., 2018). In contrast to the central nervous system, the mammalian OE retains a lifelong ability to renew neurons and rebuild functional neural circuits in response to injury. (Monti Graziadei and Graziadei, 1979; Schwob et al., 2002, 2017). Therefore, OE has been well studied as a model for neurogenesis, neural circuit formation and organ regeneration. (Talamo et al., 1989; Lavoie et al., 2017). OE consists mainly of three types of cells: basal cells (BCs), sustentacular (SUS) cells, and olfactory sensory neurons (OSNs). In mammals, each OSN expresses only one functional odorant receptor (OR) gene in a mutually exclusive manner—a phenomenon called the “one-neuron-one-receptor” rule (Serizawa et al., 2004). Besides, OSNs that express the same type of OR project their axons to a specific target glomerulus in the olfactory bulb (OB)—a phenomenon called the “one-glomerulus-one-receptor” rule (Ressler et al., 1994; Vassar et al., 1994; Takeuchi et al., 2014). Based on these two rules, mice can maintain neural circuits for olfaction throughout their lives to detect more than 100,000 different odorants using about 1,000 ORs. (Buck and Axel, 1991; Buck, 2000). Although the molecular mechanisms underlying these two rules have been extensively studied, it remains unclear how each OSN expresses only one member of the OR gene family and OSNs expressing a particular OR gene converge their axons to a specific set of glomeruli on the OB. One of the reasons that have hindered the elucidation of the molecular mechanisms regulating odor map formation is the lack of in vitro systems that allow detailed observation of olfactory sensory circuit formation.

Recent advances in stem cell technology have made it possible to reconstruct functional “mini-organs” (organoids) by recapitulating remarkable self-organization and collective behavior in vitro through three-dimensional (3D) culture of somatic stem cells and pluripotent stem cells (Eiraku et al., 2008, 2013; Sato et al., 2009). Cultured organoids exhibit various multicellular phenomena of organogenesis such as pattern formation and morphogenesis in vitro, and let us ease to obtain a large amount of tissue that contributes to biological analysis. Several reports have shown that stem cells from mouse and human adult OE can be cultured to reconstruct OSN-containing tissues (Dai et al., 2018; Peterson et al., 2019). These studies focused on Lgr5 expressing GBC as a source for spheroid formation (Chen et al., 2014; Dai et al., 2018; Peterson et al., 2019). However, the basal cells of this OE-derived spheroid rarely differentiate into OSNs and do not form tissues with a pseudostratified structure as seen in vivo. In addition, there is no comprehensive and quantitative analysis of cell heterogeneity in 3D culture of GBCs at the single cell level. To understand how the two rules, “one neuron-one receptor” and “one glomerulus-one receptor,” are achieved, we need to develop means to robustly form OE organoids from stem cells, which structurally and functionally mimic OE in vivo.

In this study, we established a novel means for forming pseudostratified OE organoids from embryonic mouse OE. Single cell RNA-sequencing (scRNA-seq) reveals that OE organoids include all OE cell types, which form a three-dimensional structure similar to that of mouse olfactory epithelium. We also demonstrated that OSNs in the OE organoids could respond to odorants in vitro, and the majority of each OSN in the OE organoids expressed only one OR, recapitulating the “one-neuron-one-receptor” rule.

## Results

### Organoid formation from the developing mouse OE stem cells

In the present study, we sought to establish novel culture systems of OE organoids, promoting OSN differentiation and the formation of pseudostratified tissues. It has been reported that by culturing adult stem cells in the OE with niche factors (NFs) such as Wnt3a, EGF, Noggin, R-spondin 1, and bFGF, OE-like structures were formed in 3D culture (Dai et al., 2018). We tested whether stem cells from developing mouse OE could form organoids in the presence of these niche factors. In the dissection, the upper part was isolated from the head of E13.5 mice (Figures 1A and 1B), then the palate was removed from the nasal epithelium (NE), the rostral half of the NE was resected (Figure 1C), and finally the OE was isolated by removing the OB (Figure 1D). Immunostaining confirms that the caudal half of the NE was the OE where Tuj1^+^ OSNs were located (Figure 1E). To generate organoids from the developing mouse OE tissues, we dissociated isolated OE tissues at E13.5 into single cells and embedded it in Matrigel (Figure 1F). By culturing the single dissociated OE cells with a culture medium containing NFs, self-organized cysts were formed and gradually increased in size as culture continued (Figure 1G). The cysts size reached a plateau and kept the size until day 30 (Figures 1G and 1H). qPCR analysis showed that Lgr5 expression was maintained from day 10 to day 30 at the same level as OE in embryonic mice (Figure 1I and 1J). Furthermore, we seeded cells under diluted conditions and observed the behavior of a single cell and confirmed that the cyst formation occurred even from a single cell (Figure 1K). These results indicate that cells derived from mouse fetal OE can spontaneously form 3D structures from a single cell when cultured with NFs.

**Figure 1.**
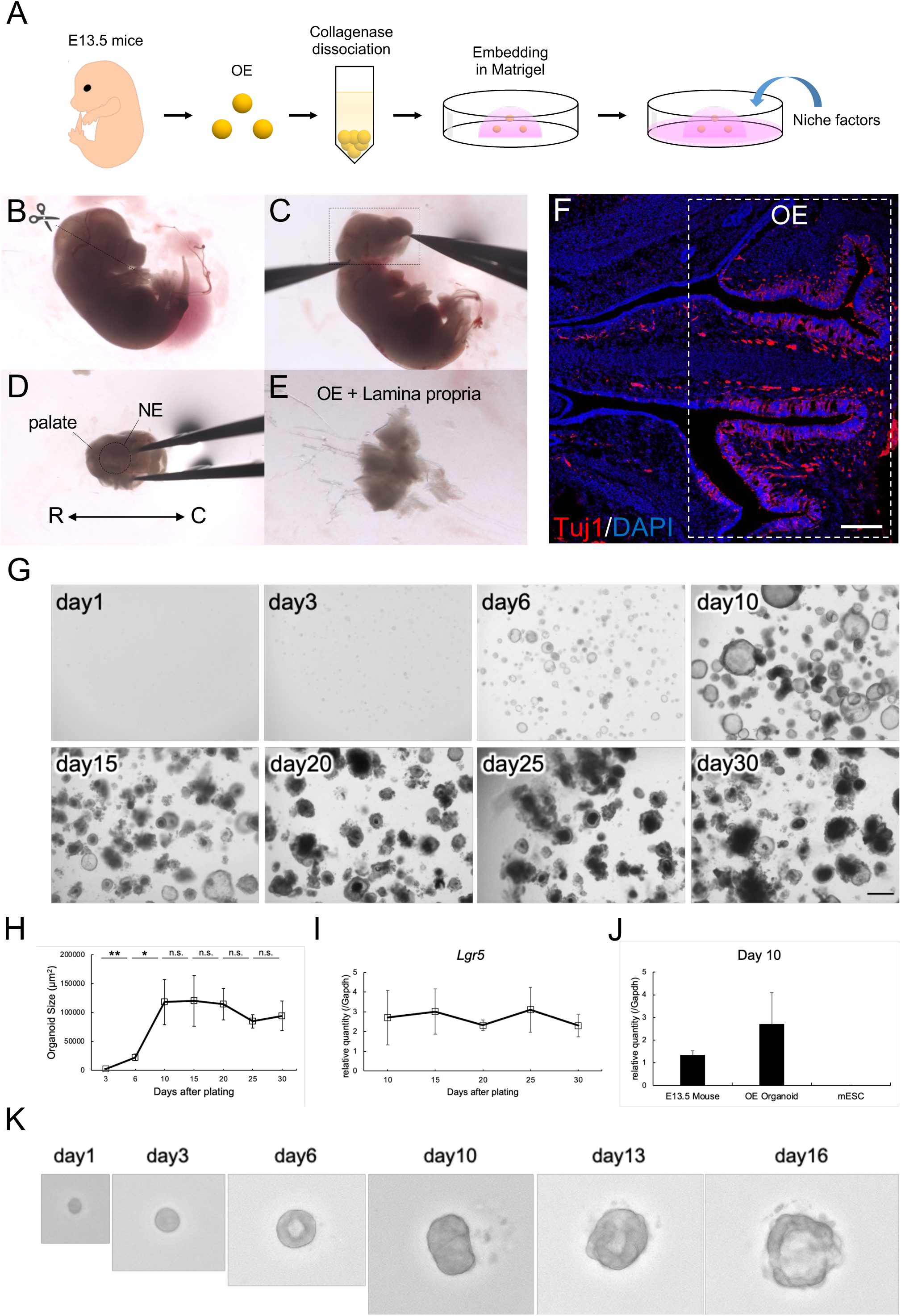
Organoid formation from embryonic mouse OE cells. (A-D) The dissection steps of mouse OE for organoid culture preparation. NE, nasal epithelium; R, rostral; C, caudal. (E) Immunostaining of E13.5 mouse nasal epithelium for Tuj1. The scale bar represents 200 µm. (F) Schematic model of the culture method for organoid formation from mouse developing OE tissues. (G) Representative bright-field images of OE organoids. Cyst-like structures are visible after 3-6 days. The scale bar represents 500 µm. (H) Temporal changes of the organoid size (n = 8). The organoids reach an area of approximately 80,000-160,000 µm^2^ within 10 days. **p < 0.01; *p < 0.05 Student’s t test. (I) qPCR for *Lgr5* in OE organoids from day 10 to 30 (mean ± s.e.m, n = 3 independent experiments). (J) qPCR for *Lgr5* in E13.5 mouse OE, OE organoids on day 10, and mESCs (mean ± s.e.m, n= 3 independent experiments). (K) Time-dependent growth of a representative cyst-like structure generated from a single cell derived from E13.5 mouse OE tissues.

### Differentiation into subtypes in OE organoids that maintain the epithelial structure

To investigate the expression of the subtype markers in the OE organoids, we performed a qPCR analysis for ΔNp63 (OE-specific HBCs marker) (Packard et al., 2011), Pax6 (SUS cells/BCs marker) (Guo et al., 2010), Mash1 (also called Ascl1; olfactory neuronal progenitor marker) (Guillemot et al., 1993), NCAM (OSNs marker), and OMP (mOSNs marker) (Lee et al., 2011) in OE organoids. ΔNp63 and Pax6 expressed at almost the same levels from day 10 to day 30, suggesting that stem cells in OE organoids were maintained at day 30 (Figures 2A and 2B). On the other hand, the expression level of Mash1 was the highest on day10 and decreased at day 30 (Figure 2C). Moreover, the expression of NCAM was maintained from day 10 to day 20, and the expression of OMP increased gradually and reached the highest levels on day 20 (Figures 2D and 2E). From day 20 to day 30, the expression level of NCAM and OMP decreased (Figures 2D and 2E). These results indicate that neuronal differentiation and maturation in the OE organoids might occur actively from day 10 to day 20, and OSNs gradually decreased after the day 20.

**Figure 2.**
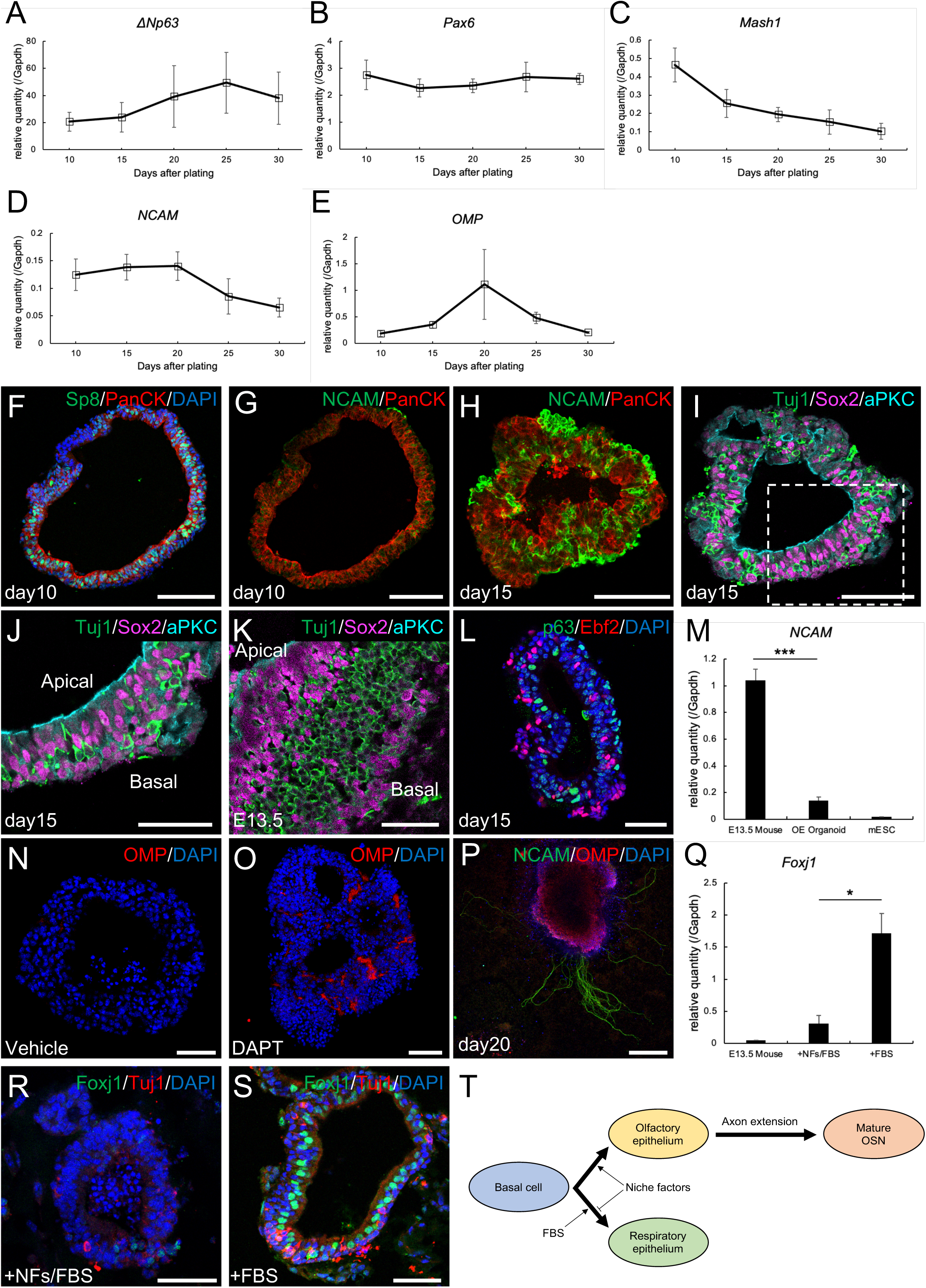
OE organoids form a polarized pseudostratified structure. (A-E) Time-course qPCR of OE organoids from day 10 to 30 (mean ± s.e.m, n = 3 independent experiments). (F-L) Cryosections of OE organoids on day 10 (F and G), on day15 (H-J and L), and E13.5 mouse OE (K). Immunostaining for Sp8 (F), PanCK (F-H), NCAM (G and H), Tuj1 (I-K), Sox2 (I-K), aPKC (I-K), p63 (L), and Ebf2 (L). (M) qPCR for *NCAM* in E13.5 mouse OE, OE organoids on day 20, and mESCs (mean ± s.e.m, n = 3 independent experiments). ***p < 0.001; Student’s t test. (N-P) The effect of Notch signaling on the expression of OMP. (N) untreated, (O) DAPT treatment (500 nM, day 10-20). (P) Quantification of the OMP^+^ cell percentages (n = 5). **p < 0.01; Student’s t test. (Q-S) The effect of NFs and FBS on the expression of Foxj1. (Q) NFs and FBS treatment (day 0-10), (R) FBS treatment (day 0-10). (S) qPCR for Foxj1 in E13.5 mouse OE, OE organoids on day 10 treated with NFs and FBS, and OE organoids on day 10 treated with FBS (mean ± s.e.m, n = 3 independent experiments). *p < 0.05; Student’s t test. (T) Schematic of olfactory/respiratory epithelial differentiation from olfactory basal cells by treatment of NFs or FBS. The scale bars represent 100 µm (A-D), 50 µm (E-G, N, O, Q, and R).

We next investigated the cell types composing the OE organoids by immunostaining. On day 10, olfactory placode markers (Sp8 and PanCK) were detected in almost all cells consisted of the cysts (Figure 2F; Kasberg et al., 2013; Comte et al., 2004; Sjödal et al., 2007). OE is mainly composed of cells of three types; the olfactory sensory neurons (OSNs), the sustentacular (SUS) cells, and the basal cells (BCs); horizontal basal cells (HBCs) and globose basal cells (GBCs) (Moulton and Beidler, 1967; Monti Graziadei and Graziadei, 1979). Basal progenitor cells reside in the basal compartment, OSNs in the middle compartment, and SUS cells in the apical side of the OE. Although very few populations of cells expressed NCAM (OSNs marker) (Carter et al., 2004) on day 10, NCAM^+^ OSNs were detected on day 15, indicating that extending culture promoted differentiation into OSNs (Figures 2G and 2H). Expression of aPKC was detected on the luminal side of the cysts on day 15, indicating that the OE organoids maintained their apical-basal polarity and formed an apical surface inside (Figures 2I and 2J). Furthermore, the expression of Sox2, a marker for SUS cells/BCs, was separated into apical and basal sides by Tuj1-positive OSN cells, indicating that OSNs, SUS cells, and BCs form a structure similar to that found in vivo. (Figures 2I-2K). We also found that p63 (a marker for HBC) is expressed in the most basal region (the outside of the cyst) (Packard et al., 2011), and Ebf2 (a marker for immature and mature OSN) is expressed in the middle region of the cyst (Wang et al., 2004) (Figure 2L). These results suggest that the organoids from the embryonic mouse OE consist of various types of OE cells and mimic the structural features of OE.

Although the expression level of neuronal markers was high on day 20, it was significantly lower than E13.5 embryonic OEs (Figure 2M). It has been reported that inhibition of the Notch signaling promote OSN production in OE (Herrick et al., 2017; Dai et al., 2018). Therefore, we next tested the treatment with gamma-secretase inhibitor DAPT (inhibiting Notch cleavage and activation) in OE organoids on day 10. Immunostaining revealed that on day 10 after DAPT administration (day 20 after in vitro culture), the ratio of OMP^+^ mOSNs was significantly increased with the treatment of DAPT (Figures 2N and 2O), suggesting that Notch inhibition promotes the maturation of OSNs of OE organoids. We also noticed that the production of OSNs was stimulated by adherent culture of OE cysts without the addition of DAPT. In the OE organoids attached on the culture dish surface, OMP expression was upregulated (Figure 2P).

### Biased differentiation into nasal respiratory epithelium (NRE) in the organoids culture with serum

In conventional neurosphere cultures, the serum has been known to affect the differentiation and maintenance of stem cells. Therefore, we next examined the effect of serum presence or absence on the OE organoids culture. we have tested three media; the medium containing NFs and 10% FBS (NFs(+)/FBS(+) medium), containing 10% FBS only (NFs(−)/FBS(+) medium), and containing neither NFs nor FBS (NFs(−)/FBS(−) medium). Cyst formation was observed in culture with all medium tested, but cells hardly grew with NFs(−)/FBS(−) medium (Figure S1A). The expression level of Lgr5, which has been reported to be involved in stem cell maintenance, was similar in all culture media. (Figure S1B). The cysts cultured with NFs/FBS and FBS expressed Sox2, an early marker for all cells of the nasal epithelium (Figures S1D and S1E) (Håglin et al., 2020). When culturing with NFs(+)/FBS(+) culture system, the growth rate of the number of cells tended to be large, indicating that both NFs and FBS promoted cyst growth (Figure S1C).

In addition to the olfactory epithelium (OE), the nasal respiratory epithelium (NRE) also develops from nasal placodes (Håglin et al., 2020). Foxj1, a marker of ciliated cells in NRE (Pardo-Saganta et al., 2015; García et al., 2019), was rarely expressed in OE organoids formed in NFs(+)/FBS(+) medium (Figure 2Q). On the other hand, we found that the cysts cultured in NFs(-)/FBS(+) media more highly expressed Foxj1 than with NFs(+)/FBS(+) (Figures 2Q-2S). Therefore, these results suggest that differentiation of OE progenitor/stem cells into nasal respiratory epithelial (RE) cells is enhanced by culturing organoids with FBS, and the differentiation is inhibited by the addition of NFs (see Figure 2T). Tuj1 was also expressed in the cysts in NFs(-)/FBS(+), indicating that the organoids consisted of both nasal RE cells and OE cells (Figure 2R).

### Single-cell RNA sequencing of OE organoids on day 10 culture

Next, single-cell RNA sequencing (scRNA-seq) was applied to OE organoids to unbiasedly investigate cell heterogeneity. We performed scRNA-seq using 10x Chromium system by preparing cells dispersed from day10 and day 20 OE organoids cultured in NFs(+)/FBS(-) media (NF-medium) and day10 OE organoids cultured in NFs(-)/FBS(+) media (FBS-medium), respectively.

For OE organoids cultured on day 10, unsupervised clustering was used to divide 11312 cells prepared from OE organoids cultured in NFs-medium and 4228 cells from OE organoids cultured in FBS-medium into 32 and 25 distinct clusters, respectively We combined several less well-defined clusters and identified 10 major cell types based on the expression of each subtype marker gene (Figure 3A and B). Consistent with the results of qPCR and immunostaining analysis, we found more Foxj1-positive cells in OE organoids induced in FBS-medium (20.9% vs. 5.8%). Furthermore, the population of NRE-type HBCs expressing Krt19, which is known to be explicitly expressed in NREs, was increased in OE organoids cultured in FBS-media. We also noticed that FBS may promote the survival and proliferation of non-OE cells: Prrx1/Pdgfra-expressing mesenchymal cells were abundant in OE organoids cultured in FBS-media, whereas these cells were almost absent in NFs-media. Conversely, cells expressing Ebf2, an OSN marker, were found in 2.6% of OE organoids cultured in NF-medium, whereas only 0.4% of cells were found in the environment, including serum (Figure 3B, 3C and 3D). These results suggest that niche factors function to maintain OE differentiation homeostasis and inhibit the differentiation and proliferation of non-OE cells.

**Figure 3.**
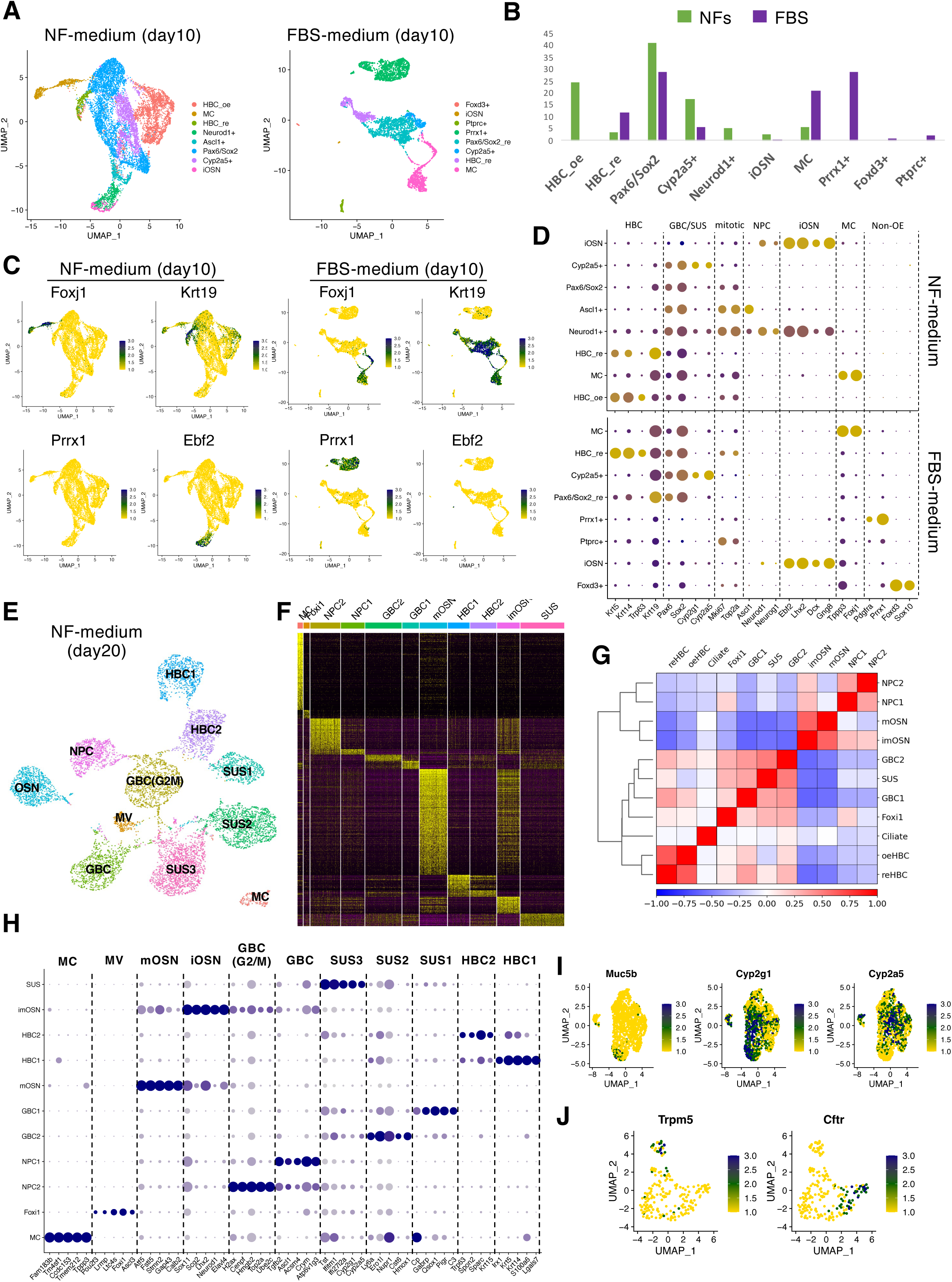
Analysis of cellular organization in OE organoids by scRNAseq. (A) Visualization of clustering results of day10 OE organoids cultured in NF- or FBS-medium on UMAP. (B) Cell number ratio in each cluster shown in A (C) Visualization of expressions of markers for NRE (Foxj1, Krt19), mesenchymal cells (Prrx1) and OSN (Ebf2) on UMAP. (D) The expression of each subtype marker in OE organoids cultured under each condition is shown in the dot plots. (E) DESC-assisted clustering result of day 20 OE organoid on UMAP. Each cluster was named according to the cell subtype identified by the gene expression pattern. (F) Genes specifically expressed in each cluster (DEGs) are shown in heatmap. (G) Heatmap of correlation coefficients between clusters using all gene expression patterns. (H) The dot plot showing the expression of each subtype marker in OE organoids on day 20. (I) The SUS cell population was re-clustered and embedded into UMAP to visualize the expression of specific marker for BG (Muc5B) and SUS (Cyp2g1, Cyp2a5). (J) The MV cell population was re-clustered and expanded into UMAP to visualize the expression of specific marker for MVs (Trpm5, Cftr).

### Single-cell RNA sequencing analysis of OE organoids on day 20 culture

Next, the cellular heterogeneity of OE organoids was investigated by scRNA-seq after further extended culture in NF-media (day 20). The scRNA-seq analysis of mouse OE-derived cells has been previously reported (Fletcher et al., 2017, Cell Stem Cell). In Fletcher et al., authors analyzed the Sox2-positive cell lineage and identified all cell types of OE. Comparing the data obtained in the current experiment with previously reported scRNA-seq data, almost all of the OE cells identified in the previous analysis were included in the our analysis. By the expression of known subtype markers, mature OSNs (Omp+/Gng13+), immature OSNs (Gng8+), neural progenitor cells (NPC; Neurod1+/Neurog1), GBC (Ascl1+), HBC (Krt5+), MV (Foxi1+), and SUS (Cyp2g1+) cells were also identified in the OE organoids at 20 days (Figure 3H). However, clustering by Seurat could not clearly separate SUS and GBC cells. This may be due to a large number of cells in this analysis (9049 cells in this study). A novel machine-learning method known as deep-embedding algorithm for single-cell clustering (DESC) was utilized to ensure accurate clustering for extensive data set through iterative learning (Li et al., 2020 Nat Comm). Therefore, we adapted a clustering method assisted by DESC. Eleven distinct cell type clusters emerged with optimal DESC resolution (0.4), and almost all cells could be accurately classified (Figure 3E). Known cell type-specific markers were used to annotate the clusters allowing the identification of HBC, SUS, microvillous cell (MV), multiciliated cell (MC), GBC, immature OSN (iOSN), and mOSN (Figure 3E and 3F). Clustering of the correlation coefficients of the expression patterns of all genes in cells excluding MCs divided them into three main groups; OSN group (GBC(G2/M)/GBC/iOSN/mOSN), SUS group (SUS1/SUS2/SUS3/MV), and HBC group (HBC1/HBC2) (Figure 3G). Both HBC1 and HBC2 express the HBC markers Krt5/Krt14/Trp63, but Irx1/Irx2 are expressed explicitly in HBC1, as revealed by the search for cluster-specific genes (Figure 3F). Cells of the SUS lineage expressing Elf5 were divided into three clusters: Pigr and Hoxm1 were identified as genes expressed explicitly in SUS1 and SUS2, respectively; SUS3 was probably the most mature SUS cell lineage, and strong expression of Cyp2g1 and Cyp2a5 was observed (Figure 3E and 3H). On the other hand, GBCs expressing Ascl1 were divided into two clusters according to the expression of cell cycle genes (GBC(G2/M) and GBC). Muc5B, a marker of Bowman’s gland cells, was expressed in some cells of the SUS3 cluster. Furthermore, re-clustering of MV cells revealed that two types of cells, one specifically expressing Trpm5 and the other specifically expressing Cftr, were differentiated in OE organoids on day 20. These results suggest that organoids formed from the OE of E13.5 mouse embryos reconstitute almost all cell types contained in the OE.

### Lineage trajectory analysis and identification of pseudotime-dependent genes in OE organoids

We next attempted to use the scRNA-seq data to infer the lineage trajectory of stem cells in OE organoids. In the PCA plot of scRNA-seq data obtained from OE organoids of day20, cells are arranged in a bifurcated fashion, with cells expressing HBC marker (Krt5), OSN marker (Ebf2), and SUS marker (Cyp2g1) plotted at the three endpoints, respectively (Figure 4A). These results suggest two major lineage trajectories from HBC to OSN and SUS in OE organoids. In the previous reports, it has been suggested that there are three branching lineage trajectories from HBC to SUS, MV, and OSN (Fletcher et al., 2017, Street et al., 2018). Therefore, we applied slingshot, which are algorithms for inferring cell lineage trajectory, to scRNA-seq data from OE organoids on day 20 to investigate whether the same stem cell homeostasis is reproduced in OE organoids as in vivo. As in the previous report using OE, three trajectories were inferred: from HBCs to SUS and from HBCs to MVs or OSNs via GBCs (Figures 4B, 4C and 4D).

**Figure 4.**
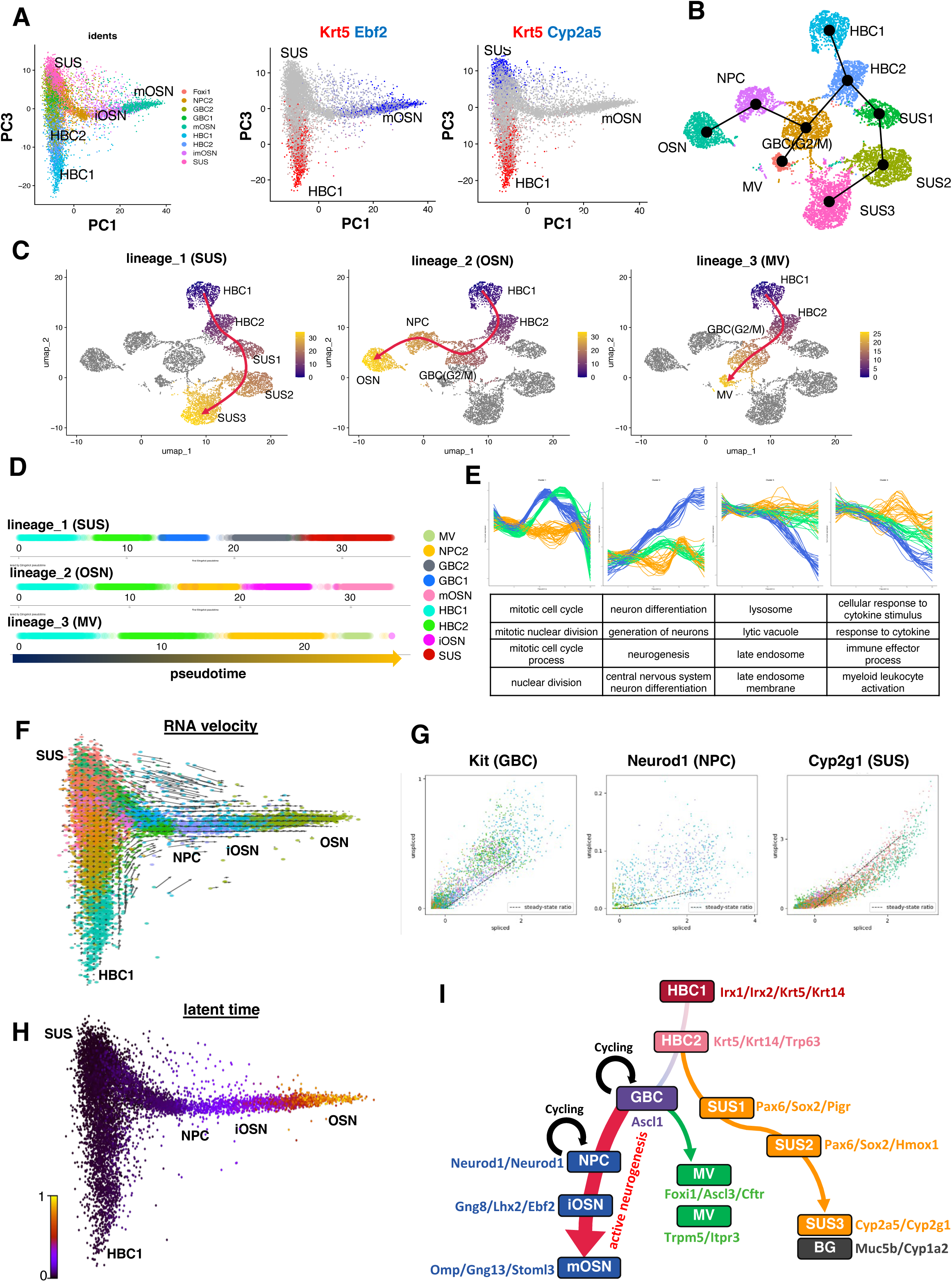
Lineage trajectory and RNA velocity of OE organoid on day 20. (A) Dimensionality reduction of scRNA-seq results using first and third principal components. The left panel shows color coding by DESC-assisted clustering result. The middle and right panel show the expression of Krt5 (red), Ebf2 (blue, in middle panel) and Cyp2a5 (blue, in right panel) (B) Inferred lineage trajectory by Slingshot. (C) Visualization of pseudotime of 3 different trajectories estimated by Slingshot on UMAP. (D) The result of arranging the cells of each trajectory on a line along the pseudotime. (E) Normalized gene expression along pseudotime of the top four clusters of highly variable genes extracted by Tradeseq. The bottom panel shows the results of enrichment analysis of the genes included in each cluster. (F) RNA vectors of day20 OE organoids were calculated with scVelo and overlayed on the PCA plot (PC1/PC3). (G) The phase portraits of marker genes. (H) Calculated latent time by scVelo was overlayed on the PCA plot (PC1/PC3). (I) The proposed model for cell lineage relationships during OE organoids formation.

We then applied Tradeseq (optimized number of nknot was nine), which can detect trajectory-dependent gene expression changes, to the data. After fitting the negative binomial generalized additive model (NB-GAM) with the fitGAM function, the results of comparing the expression of marker genes along the pseudotime between lineages show that the estimation of lineage trajectory by slingshot is reasonable (Figure 4E). We also identified trajectory-dependent genes, and these genes were classified into 33 clusters according to their pseudotime-dependent expression patterns (Figure 4F). The cluster containing the largest number of genes showed a specific upregulation during MV and OSN differentiation. Gene enrichment analysis of these genes indicates that they are related to the cell cycle. On the other hand, these gene sets were hardly expressed in the SUS lineage trajectory, suggesting that, unlike GBC-mediated cell differentiation in OSN and MV, the pathway from HCB to SUS is a direct differentiation pathway that does not associate with cell division. A second cluster is a group of gradually upregulated genes along the OSN lineage and has functions mainly related to neural differentiation (Figure 4F). The fourth cluster, which is transiently up-regulated in the SUS lineage trajectory, is composed of genes involved in inflammatory responses. To investigate the dynamics of differentiation transition in OE organoids, we analyzed RNA velocity using scVelo, that describes the rate of gene expression change based on the ratio of its spliced and unspliced messenger RNA (Bergen et al., 2020 Nat Biotech). When RNA velocities derived for OE organoids on day 20 are projected into a PCA plot of cells clustered by Seurat, we found a strong velocity pattern originating from a proliferating GBC, and proceeding through a sequence of NPC to a more mature differentiated OSN (Figures 4F and 4G). The remaining cell population show no clear RNA velocity flow. The phase portraits of marker genes suggests that Kit and Neurod1 expression is upregulated in GBC and NPC populations, while Cyp2g1 and Ascl3 are in steady state or decreased. These results indicate that OSN is actively produced through neurogenesis from GBC in the OE organoid on day20, whereas HBC, SUS and MV are either differentiated or arrested stem cell state.

Taken together, a series of analyses for scRNA-seq data revealed that organoid cultured from embryonic mouse OE showed similar stem cell homeostasis to that *in vivo* and that the neurogenesis of OSNs continued actively, while the production of SUS and MVs was almost complete until culture day 20 (Figure 4H).

### The functionality of OSN in OE Organoid

Next, we examined the functional maturation of OSNs in OE organoids. During OE development, the OSNs extend axons to the OB and transmit odorant information to the brain. To verify the axon growth ability of OSNs induced in OE organoid cultures, we cultured the OE organoids in a more suitable condition for OSN maturation, in which OE organoids were transferred to glass-bottom dishes on day 10 and embedded in collagen gel (Figure 5A). After 10 days under these conditions, axon elongation from OE organoids was observed. Immunostaining showed that the axons were positive for NCAM and OMP (Figure 5A). Next, Ca2+ imaging was performed on the OE organoids with elongated axons. When a saline solution was added, little neuronal activity (Ca2+ surge) was detected. However, when isovaleric acid (IVA), one of the odorants, was added (Kim et al., 2018), spontaneous Ca2+ oscillations were observed, suggesting that the OSNs of the OE organoids responded to the odorant in vitro, suggesting that the OE organoid OSNs reacted with odorants in vitro (Figure 5B).

**Figure 5.**
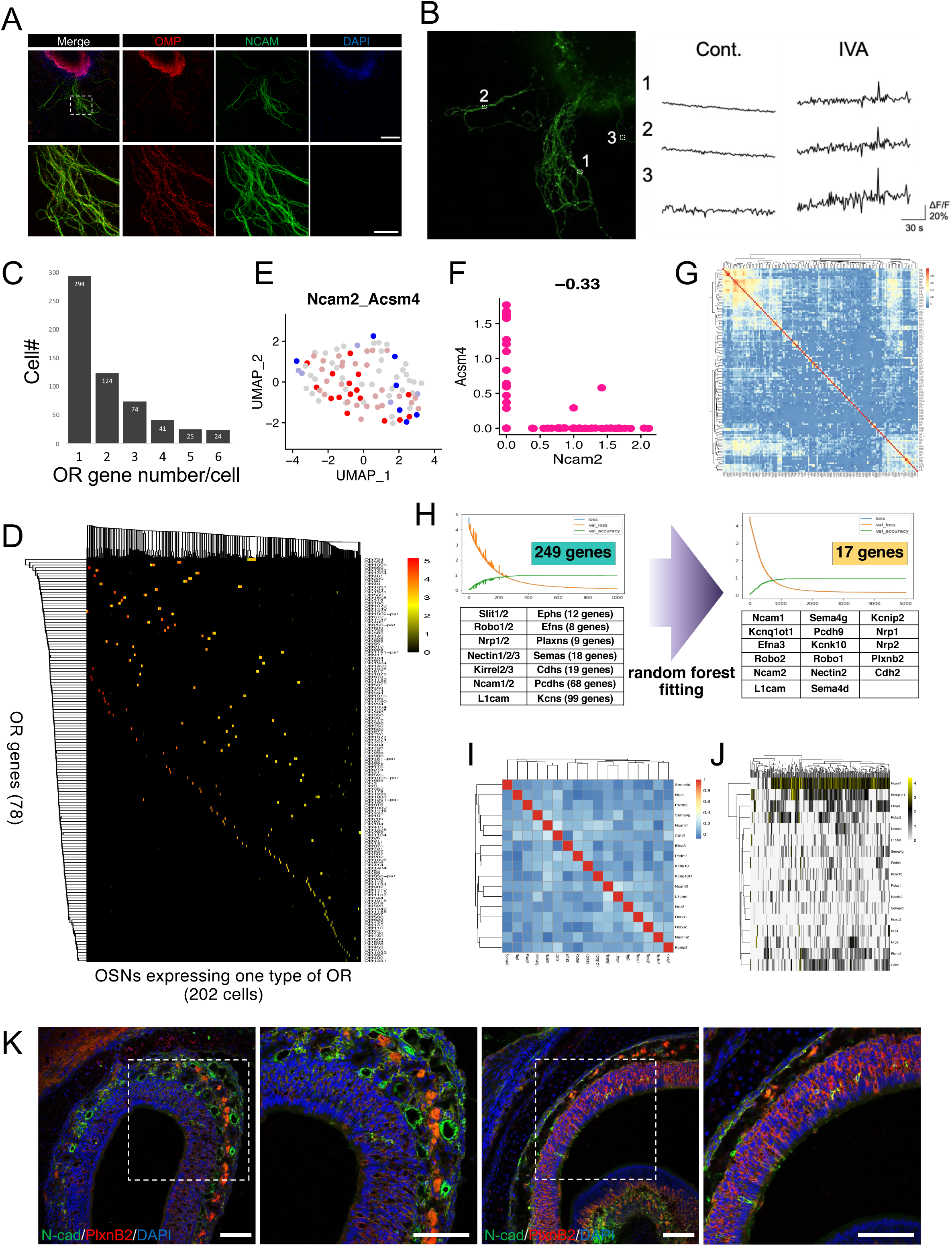
Analysis of the functionality of mature OSNs and identification of molecular codes for classifying OSNs by OR expression. (A) Immunostaining for axons extended from OE organoids on day 20. The scale bars represent 200 µm (upper panel) and 50 µm (lower panel). (B) Analysis of Ca^2+^ dynamics in individual axons on day 20. Numbers correspond to individual ROIs analyzed simultaneously by Ca^2+^ imaging. (C) Number of OR genes expressed in OSNs of OE organoids on day 20. (D) Heatmap showing the expression of each OR gene in OSNs that exclusively express one type of OR. (E) The matured OSN was re-clustered and embedded into UMAP to visualize the expression of Ncam2 (blue) and Acsm4 (red). (F) Scatter plots based on the expression of Ncam2 and Ascm4. (G) Heatmap showing the correlation coefficients of expression of 249 axon guidance-related genes in mature OSNs. (H) Discrimination of OSNs based on OR classes using a deep neural network. The learning curve using 249 genes is shown in the left panel, and the learning curve using the remaining 17 genes after feature reduction by random forest fitting is shown in the right panel. (I) Heatmap showing the correlation coefficients of expression of the remaining 17 genes in mature OSNs (J) Heatmap showing the expression of the remaining 17 genes in mature OSNs. (K) Immunostaining for Cdh2 and Plxnb2 in mouse OE.

We next examined the repertoire of odorant receptor molecules expressed in OSNs using scRNA-seq data. First, we extracted mature OSNs highly expressing Omp from OE organoid cells on day 20 (599 cells; normalized expression level of Omp >1 in scRNA-seq data). We found that 294 OSNs (49.0%) exclusively expressed only one type of OR gene. On the other hand, 124 OSNs (20.7%) expressed two types of ORs, and 181 OSNs (30.2%) expressed three or more ORs simultaneously (Figure 5C). Of the 294 cells selectively expressing one type of OR gene, 202 cells showed exceptionally high OR gene expression levels, which were distinguished by 78 different OR genes (Figure 5D). These results indicate that the “one-neuron-one-receptor” rule is at least partially achieved by an intrinsic mechanism in OSNs, even in cultured organoids without specific external stimuli and projection targets.

Recent studies have shown ORs themselves regulate the transcriptional levels of molecules involved in the odor-map formation, such as axon guidance molecules and cell adhesion factors. (Imai, Sakano, 2006; Sakano, 2010). Ncam2 (Ocam) and Acsm4 (Omacs), which show mutually exclusive expression patterns in OSNs, were also expressed mutually exclusively in OSNs of OE organoids (Figures 5E and 5F), implying that not only the exclusive selection of ORs but also the associated transcriptional regulation of axon guidance molecules may be recapitulated in the organoid culture. Taking advantage of these phenomena of OE organoids, we next attempted to screen for informative molecules for OSN classification for ORs. First, we examined the expression of 249 axon guidance-related genes in 202 OSNs, distinguished by 78 different ORs (Figure 5). The correlation matrix shows that some of these genes show similar expression patterns in each OSN, while others differ depending on the type of OSN (Figure 5G). Using the expression patterns of these genes, we trained a classifier to discriminate 78 classes of OSNs by a deep neural network under Tensorflow and KRAS framework and obtained a classifier with almost 100% probability of a correct rate within 200 epochs (Figure 5H). Using this model, we reduced the redundancy of features by random forest fitting and finally identified 17 genes as the minimum number of combinations that can be used to discriminate 78 classes of OR-expressing OSN cells (Figure 5H). Among the 17 genes, there were 12 genes (Ncam1, Ncam2, Pcdh9, Nrp1, Nrp2, Robo1, Robo2, Efna3, Kcnq10, L1cam, Nectin2, Sema4d) whose expression and function in OSN have been previously mentioned, while 5 genes (Sema4g, Kcnip2, Kcnq1ot1, Plxnb2, Cdh2) that have never been mentioned with OSN were also identified. We confirmed by immunostaining the expression patterns in mouse OE of Plxnb2 and Cad2, which we newly identified as molecules with variable expression in OSNs. Cdh2 was expressed in a small fraction of OSN (Figure 5L). On the other hand, Plxnb2 shows a region-specific expression in OSN. Plxnb2 was expressed in most OSNs in the posterior OE, whereas it was almost absent in the anterior OE (Figures 5L and S2). These results demonstrate the usefulness of OE organoids as a model to study the mechanism of selective OR expression and odor map formation in OSNs.

## Discussion

In this study, we show the formation of organoids that recapitulate the olfactory epithelium development. The organoid mainly contains basal cells, sustentacular cells, and olfactory neurons.By culturing cells from embryonic mouse olfactory epithelium (OE) tissues in the presence of specific niche factors, a pseudostratified OE-like structure can be stably induced, mimicking the structure of the mouse OE. The previously reported methods for tissue formation from adult mouse OE stem cells are not adequate as a model of the olfactory system. This is because they cannot consistently form polarized pseudostratified structures and have low efficiency in inducing OSNs (Dai et al., 2018). In the current study, one of the improvements is the generation of organoids using cells from the embryonic OE at mouse E13.5. This is the stage when the OE begins to be divided into three layers by immature OSNs, and a more persistent wave of neuronal differentiation takes place (Cau et al., 2002; Ikeda et al., 2007; Sokpor et al., 2018). As a result, stem cells in the embryonic OE may form epithelial tissues with more robust polarized structures and differentiate more actively into neurons compared to adult OE stem cells.

Consistent with previous reports, embryonic OE stem cells also required the addition of niche factors to maintain basal cells in OE organoids. We found that a culture environment with FBS instead of niche factors promoted a biased induction of respiratory epithelial (RE) cells (Foxj1+) at the expense of differentiation into neurogenic lineages. Both OE and RE horizontal basal cells (HBCs) are known to act as stem cells for regeneration in response to injury (Leung et al., 2007; García et al., 2019). Growth factors such as EGF and FGF in fetal bovine serum (FBS) may encourage the transformation of OE stem cells into RE cells. However, it’s also possible that these factors promote the growth and differentiation of a small number of contaminated RE stem cells. More research is required to comprehend the characteristics of nasal epithelial stem cells for transdifferentiation.

scRNA-seq analysis revealed that the OE organoids on day 20 contain all the cellular lineages of OE. HBCs expressing Krt5, Krt14 and Trp63. were divided into two populations (HCB1/HCB2). HBC1, which exhibits higher expression levels of Krt5 and Trp63, is considered to be a more undifferentiated population of HBC. We identified Irx1/2 as molecules that are specifically expressed in HBC1. Consistent with previous reports, inference of cell lineage trajectory from scRNA-seq data suggested two major different cell trajectories originating from HBC in OE organoids; a direct pathway from HBC to SUS without cell division, and a pathway from HBC to OSN or MV via proliferating GBCs. We also noticed that there are two cell types of MV cells expressing Trpm5 or Cftr, that is tuft cell and ionocyte, respectively. The analysis of RNA velocity showed that OSN was actively produced in OE organoids on day 20, while SUS and MV production was nearly complete. This finding aligns with the fact that the expression of cell cycle markers is mainly limited to neural lineage cells, GBCs, and NPCs. These results indicate that similar mechanisms of cell differentiation and tissue homeostasis may be occurring in organoid cultures as in living organisms. One current limitation is that it is unknown from which cell types in the OE the organoids are formed. From the analysis of marker expression patterns, it is considered that niche factors maintain proliferative ability of basal cells, but it remains unclear which GBCs or HBCs are involved in organoid formation. Although it has been reported that niche factors maintained Lgr5^+^ GBCs (Dai et al., 2018), our data have shown that HBCs were also maintained. Therefore, it is assumed that HBCs might express Lgr5 and the Lgr5^+^ HBCs might be maintained by niche factors. Otherwise, niche factors might act as some kind of signaling factor to maintain the proliferative potential of HBCs. It is necessary to investigate whether similar organoids can be formed using basal cells, such as GBCs and HBCs, only.

In our culture system, we found that inhibiting Notch and promoting axon extension in organoids can help mature olfactory sensory neurons (OSNs). Previous research has shown that OSNs mature and express Omp after their axons extend to the olfactory bulb (Liberia et al., 2019). Therefore, organoids with extended axons may mimic the natural process of OSN maturation. We also observed that OSNs in these organoids responded to odors in vitro using calcium imaging. Studying how mice detect different odors using their approximately 1,000 ORs in various combinations is challenging. On the other hand, OSNs in organoids express a limited number of ORs, making it easier to study their response to odors. Thus, the organoids could be useful for investigating the function of ORs in future studies.

It’s important to note that about half of the olfactory sensory neurons in OE organoids express only one type of olfactory receptor, following the “one-neuron-one-receptor” rules. Recently, there has been active research on how OSNs express only one OR gene in a monoallelic and stochastic fashion using mouse models. OR expression is regulated by both “positive regulation” and “negative regulation” (Serizawa et al., 2004). Markenscoff-Papadimitriou et al. demonstrated that OR expression is positively regulated by interchromosomal interactions (Markenscoff-Papadimitriou et al., 2014). Our data has shown that OE organoids express a limited variety of ORs. With this in mind, examining the epigenetic states of OE organoids, could be useful in understanding the positive regulation of OR gene selection. Unlike positive regulation, little is known about negative regulation. Roughly half of the OSNs in OE organoids expressed multiple ORs. During development, OSNs express multiple ORs, but as they mature, they eventually choose only one type of OR (Tan et al., 2015). This means that OE organoids can be used to study how OSNs narrow their OR expression to just one during maturation. By examining the OR profiles of OSNs in OE organoids created from a single cell, we can also gain insight into the stage of development when genomic regulation allows for the exclusive selection of a single OR. In addition to the OE organoid, creating an in vitro model that incorporates the olfactory bulb will enhance our understanding of the molecular mechanisms involved in forming the neural circuitry of the olfactory system in the future. If it is possible to generate OE from pluripotent stem cells, such as mouse ES cells, using current culture conditions, it will be easier to obtain large quantities of cell samples. This will enable multimodal omics analysis and CRISPR screening, which require a large number of cells. If this is achievable, it is hoped that this experimental system will contribute to a comprehensive understanding of the olfactory system.

## Method details

### Mouse OE Organoid Culture

Single cells from OE were cultured according to previously reported protocol with some modification (Sato and Clevers, 2013; Drost et al., 2016). We prepared pregnant female mice and dissected OE of E13.5 ICR mouse embryo in the PBS at room temperature. After dissecting, OE was digested in 0.5% collagenase Type I (Worthington) in PBS with 10 µM Y-27632 for 1 hour at 37℃. Cells were dissociated gently by pipetting for about 10 times. After adding 10 ml of DMEM/F12/Glutamax-1 (Gibco) to wash, cells were centrifuged at 1,000 rpm for 3 minutes. The cell pellet was resuspended in 1 ml of TrypLE (Gibco) with 10 µM Y-27632, and then digested for 15 minutes, with pipetting up and down with a P1000 pipette every 5 minutes. After adding 10 ml of DMEM/F12/Glutamax-1 (Gibco) to wash, cells were counted using hemocytometer and centrifuged at 1,000 rpm for 3 minutes. After resuspending in Matrigel, 5,000 cells in a 10-µl drop in the middle of one well of a 48 well plate (FALCON). The 48 well plate was plated upside down in the CO_2_ incubator (5% CO_2_, 37 ℃) for about 15 minutes. The cells were cultured in 250 µl of conditioned OE organoid growth medium which is optimal for the culture of Lgr5^+^ organoids according to the previous report with some modification (Ren et al., 2014; Dai et al., 2018). The ingredients of growth medium are based on DMEM/F12/Glutamax-1 (Gibco) supplemented with 200 ng/ml R-spondin 1 (R&D), 100 ng/ml Noggin (R&D), 50 ng/ml Wnt3a (R&D), 50 ng/ml EGF (PeproTech), 20 ng/ml bFGF (Wako), 1% N2 supplement (Gibco), 2% B27 (Gibco), 1mM HEPES (ThermoFisher), 10 µM Y-27632 (Calbiochem), and 4% (v/v) Matrigel (growth-factor-reduced; BD Biosciences). Medium was refreshed every 2-3 days with medium that did not contain Y-27632. NFs(−)/FBS(+) medium is based on DMEM/F12/Glutamax-1 supplemented with 10% FBS (JRH), 1% N2 supplement, and 1mM HEPES. For forced neuronal differentiation, DAPT (500 nM) was added into OE organoids at day 10. Organoids were treated with DAPT or DMSO (used as control) for 10 days and then subjected to further analysis.

For induction of axon extension, OE organoids were transferred to glass bottom dish and embedded in collagen gel (Cellmatrix Type I-A; Nitta Gelatin) at day 10. Organoids were cultured with OE organoid growth medium for 10 days. A one-half volume of the medium was changed every three days.

### OE Dissection from Mouse Embryos

OE was dissected from E13.5 mouse embryos according to previously reported protocol with some modification (Fouquet and Trembleau; 2019). The upper part of the head was isolated by cutting through the mouth on both sides and the head was inverted to see the underside. The lateral sides of the developing palate in the horizontal plane were cut and a coronal section of the tissue through the septum and the turbinate bones was made. The caudal part of the nasal cavity was detached from the OB and the lateral side of the turbinate was cut.

### Quantitative PCR

We used organoids in 1-2 wells of 48 well plate and 3 of amputated-mouse OE in each qPCR analysis. Total RNA purified from samples using the RNeasy micro kit (QIAGEN) and cDNA synthesis was performed by using SuperScript II (Invitrogen). qRT-PCR was performed with Power SYBR Green PCR Master Mix by the QuantStudio 6 and 7 Flex Real-Time PCR Systems (Applied Biosystems) and data was calculated by standard curve method and normalized to *Gapdh* expression. See Table for primer sequences.

### Immunostaining

For tissue frozen section, OE organoids and mouse OE tissues are fixed with 4% paraformaldehyde (PFA) in PBS for 15 minutes at room temperature. Samples were then washed in PBS, immersed in 18% sucrose in PBS at 4℃ overnight, and frozen in OCT embedding compound. Sections were cut (10 µm) and permeabilized with 0.3% Triton-X 100 in PBS for 15 minutes, and then blocked for 15 minutes at room temperature in blocking buffer (2% skim milk in PBS containing 0.3% Triton-X 100). For first antibody incubation, antibodies were diluted in blocking buffer and incubated overnight at 4℃. Anti-bodies against the following proteins were used at the indicated dilutions: Sp8 (goat, 1/100; Santa Cruz), PanCK (mouse, 1/250; Sigma), NCAM (rabbit, 1/500; Chemicon), Tuj1 (mouse, 1/500; Covance and rabbit, 1/500; Covance), Sox2 (goat, 1/250; Santa Cruz), aPKC (rabbit, 1/100; Santa Cruz), p63 (mouse, 1/500; Santa Cruz), Ebf2 (sheep, 1/200; R&D), OMP (mouse, 1/400; Santa Cruz), Foxj1 (mouse, 1/1000; eBioscience), N-cadherin (mouse, 1/500; BD), PlexinB2 (rabbit, 1/25; PGI). After washes with 0.05% Tween/PBS three times for 5 minutes, samples were incubated for 1 hour at room temperature with DAPI (Nacalai) as a counter staining and appropriate secondary antibodies conjugated with the fluorescent probes, and then washed with 0.05% Tween/PBS for three times for 5 minutes. Stained sections were imaged with a confocal microscope (TCS SP8; Leica). Morphological analysis was done with image J (version 1.51, NIH). For whole mount immunofluorescence, tissues were fixed with 4% PFA in PBS for 20 minutes at room temperature. Samples were then washed in PBS, permeabilized 0.3% Triton-X 100 in PBS for 30 minutes at room temperature, and then blocked for 1 hour at room temperature in blocking buffer. For first antibody incubation, antibodies were diluted in blocking buffer and incubated overnight at 4 ℃, washed 0.05% Tween/PBS. Second antibodies were diluted in blocking buffer and incubated for 2 hours at room temperature. Stained samples were analyzed with a confocal microscope (TCS SP8; Leica).

### Ca^2+^ Imaging

For Ca^2+^ imaging, OE organoids were subjected to collagen gel embedded culture to induce axon extension in OE organoid growth medium on day 10. For calcium dye loading, OE organoids were incubated in fluo8-AM solution as described previously (Ikegaya et al., 2005). Imaging was carried out at 37℃ using a confocal microscope (CV1000; Yokogawa). Fluo8 fluorescence images were viewed through dry objective (10x). The fluorescence change over time is defined as ΔF/F = (F − F_basal_)/F_basal_, where F is the fluorescence at any time point, and F_basal_ is the fluorescence at the start of the measurement for each ROI. For pharmacological experiment, IVA (100 µM) was applied.

### Single-cell mouse OE organoid preparation for scRNA-seq

Mouse OE organoids cultured in Matrigel domes (on day 10) and in collagen gel (on day 20) were dissociated into single cells using TrypLE (Gibco) supplemented with 150 µg/ml DNase I (Roche) at 37 ℃ for 5 minutes, and then neutralized with 0.04% BSA/PBS. The dissociated organoids were pelleted and resuspend with 0.04% BSA/PBS. The resuspended organoids were then placed through 35-µm filter to obtain a single cell suspension. The single-cell suspension was proceeded with the Chromium Single Cell 3’ Reagent Kit v3.1 (10x Genomics) using a 10x Genomics Chromium Controller. A total 10,000 cells were loaded into each channel of Chromium Next GEM Chip B. A total of 10,000 cells and Master Mix were loaded into each channel of the cartridge to generate the droplets on the Chromium Controller. Beads-in-emulsion (GEMs) were transferred and GEMs¥-RT was undertaken in droplets by PCR incubation. GEMs were then broken and pooled fractions were recovered. After purification of first-strand complementary DNA (cDNA) from the post GEM-RT reaction mixture, barcoded, full-length cDNA was amplified via PCR to generate sufficient mass for library construction. Enzymatic fragmentation and size selection were used to optimize the cDNA amplicon size. TruSeq Read 1 (read 1 primer sequence) is added to the molecules during GEM incubation. P5, P7, a sample index, and TruSeq Read 2 (read 2 primer sequence) are added via End Repair, A-tailing, Adaptor Ligation, and PCR. The final libraries were assessed by an Agilent Technology 2100 Bioanalyzer and sequenced on an Illumina NovaSeq 6000 sequencer with a paired-end 100 cycle kit (28+8+91).

### Secondary analysis of scRNA-seq data

Cell Ranger 5.0.1 was used to process the raw data from the NovaSeq 6000. Next, we analyzed the processed data using Seurat v.4. Briefly, the cells with aberrantly high or low numbers of unique molecular identifiers (UMIs) and high mitochondrial gene detection rates were removed. Gene expression levels were normalized to the total number of UMIs Then, the Top 2000 highly variable genes were found by FindVariableFeature function R package Seurat, and the ScaleData function R package Seurat was used to scale all genes, and the RunPCA function was used to reduce the dimensionality of the Top 2000 highly variable genes selected. The “FindNeighbors” and “FindCluster” functions (resolution = 2) R package Seurat are used to cluster cells when dim = 20. Next, we use the RunUMAP method for further dimensionality reduction.

For day 20 sample, we clustered cells into distinct cell types using DESC (louvain_resolution =0.4), a deep learning based clustering algorithm (Li et al., Nat Comm, 2020). A gene expression matrix was generated by Cell Ranger 4.0.0 (10x Genomics) with the GRCm38 (Ensemble release 101) as a reference genome. Using the Velocyto (version 0.17) counting pipeline with Python 3.8.5 to obtain two matrices data. The data were estimated RNA velocity and latent time using Seurat version 3.9.9 in R version 3.6.3 and scVelo version 0.2.2 in Python 3.8.5, and then the selected genes were in the phase portrait. For RNA velocity analysis, loom files were generated using the Velocyto software (version 0.17). RNA velocity analysis was conducted using scVelo on Python 3.9.5. The data were estimated RNA velocity and latent time using Seurat version 3.9.9 in R version 3.6.3 and scVelo version 0.2.2 in Python 3.8.5, and then the selected genes were in the phase portrait. For trajectory analysis, the clustered data was processed using slingshot v.2 and tradeSeq v.1.

### Classification of OSN classes by deep neural network (DNN)

First, 202 OSNs were selected from the scRNAseq data at day 20 of culture, expressing only one OR gene at high levels. These cells were categorized into 78 classes based on the type of OR they expressed. 150 cells were used to build a training model using the Python library “chainer”. The model was validated using the remaining cells. Using this model, we reduced the redundancy of features by random forest fitting using “RandomForestRegressor” from sklearn module and finally identified 17 genes as the minimum number of combinations that can be used to discriminate 78 classes of OR- expressing OSN cells

## Acknowledgement

We thank Masatoshi Ohgushi and Yusuke Seto for invaluable comments. We would like to thank members of Core Facility of WPI-ASHBi for consultation and processing of transcriptome sequencing. This work was supported by Grant-in-Aid for Scientific Research (B) (Ministry of Education, Culture, Sports, Science, and Technology (MEXT), Japan (23H02487 to M.E.), Grant-in-Aid for Transformative Research Areas (Ministry of Education, Culture, Sports, Science, and Technology (MEXT), Japan (23H04933 to M.E.) and Core Research for Evolutional Science and Technology (CREST, JST) (JPMJCR12W2 to M.E.).

